# Neutrophil Extracellular Traps have DNAzyme activity that drives bactericidal potential

**DOI:** 10.1101/2023.10.23.563618

**Authors:** Ti-Hsuan Ku, Nikhil Ram-Mohan, Elizabeth J Zudock, Ryuichiro Abe, Samuel Yang

## Abstract

The mechanisms of bacterial killing by neutrophil extracellular traps (NETs) are unclear. DNA, the largest component of NETs is believed to merely be a scaffold with minimal antimicrobial activity through the charge of the backbone. Here, we report that NETs DNA is beyond a scaffold and produces hydroxyl free radicals through the spatially concentrated G-quadruplex/hemin DNAzyme complexes, driving bactericidal effects. Immunofluorescence staining showed colocalization of G-quadruplex and hemin in extruded NETs DNA, and Amplex UltraRed assay portrayed its peroxidase activity. Proximity labeling of bacteria revealed localized concentration of radicals resulting from NETs bacterial trapping. *Ex vivo* bactericidal assays revealed that G-quadruplex/hemin DNAzyme is the primary driver of bactericidal activity in NETs. NETs are DNAzymes that may have important biological consequences.

**One-Sentence Summary:** G-quadruplex/hemin DNAzymes may be major contributors to biological consequences of neutrophil extracellular traps.

## Main Text

Neutrophils represent a substantial component of the innate immune system, serving as the primary defense against infecting pathogens. Their ability to eliminate pathogens is carried out through various mechanisms, such as phagocytosis, degranulation, and the synthesis of neutrophil extracellular traps (NETs), which were discovered in 2004 (*1*). NETs are intricate networks composed of decondensed chromatin DNA fibers that ensnare and immobilize invading pathogens, effectively impeding their dissemination. Upon entrapment, the immobilized pathogens become exposed to potent and often lethal concentrations of histones, azurophilic granules, specific granules, tertiary granules, and cytosolic proteins intricately bound to the DNA fibers released during the process of NETosis. However, depending on the pathogen, like *Pseudomonas aeruginosa* and *Staphylococcus aureus*, NETs may have a bacteriostatic effect rather than bactericidal and eventually aid with killing through the complement (*2*, *3*). Additionally, conditions inducing NETosis also affect the NETs bactericidal properties (*2*). Although NETs play a significant role in eliminating pathogens, they can also contribute to various immunopathologies, such as delayed tissue repair, inflammation, vaso-occlusion, and autoantibody generation (*4*). Even though the individual constituents of NETs are known to be involved in both pathogen clearance and the development of immunopathologies, the overall mechanism by which NETs operate remains incompletely understood. Despite being considered a mere scaffold for associated proteins and pathogen sequestration, DNA, the most prevalent component of NETs, has recently been found to play a more active role. This is evidenced by the observation that DNase treatment leads to a two-fold increase in bacteremia, abolishment of NET-mediated cytotoxicity *in vitro*, reduction in levels of citrullinated histones associated with NETs, and direct disruptment of bacterial cell membrane through cation chelation (*5*).

DNA has the capacity to adopt diverse secondary structures beyond the conventional B-form, including the i-motifs, triple helices, and G-quadruplexes (G4). Through Hoogsteen hydrogen bonding, four guanine bases can come together to create a square planar structure known as a guanine tetrad (G-tetrad or G-quartet), and the stacking of two or more G-tetrads results in the formation of a G4 structure. The utilization of genome-wide high-resolution sequencing revealed the presence of over 700,000 unique G4 structures within the human genome, with a notable enrichment observed in promoter regions surrounding transcription start sites (*6–8*). Recent physiological findings have established a noteworthy interplay between G4 structures and hemin within living cells, wherein G4 structures exhibit a remarkable capacity to sequester free hemin, thereby serving as a protective mechanism against hemin-induced oxidative damage (*9–11*). The G4/hemin (G4/H) complex has been recognized for its ability to emulate the functionality of a peroxidase, facilitating the breakdown of hydrogen peroxide (H_2_O_2_) to generate hydroxyl free radicals (OH•) (*12*). This process can occur directly or via superoxide intermediates, ultimately driving the Haber-Weiss reaction (*13*, *14*). While the DNAzyme activity of the G4/H complex has been well-established through various *in vitro* studies, and the presence of G4 structures in the genome and their role in sequestering hemin have been demonstrated *in vivo*, it remains unclear whether G4 structures exist in the extruded DNA of NETs, their capacity to form the DNAzyme with hemin and generate OH•, and their contribution to bactericidal activity in NETs.

### Co-localization of G4 and Hemin in NETs

To determine if G4 and hemin are endogenously present within neutrophils and extruded NETs, we performed *in vitro* immunofluorescence co-localization of G4 and hemin on protein-bound crude NETs. NETosis was stimulated using *Escherichia coli* in negatively selected neutrophils from healthy donors. G4 and hemin were subsequently labeled with BG4 antibody and anti-hemin monoclonal antibodies respectively. NETs formation was confirmed by labeling with anti-citrullinated histone antibody and subsequently examined using bright field and fluorescent microscopy (Fig. 1). Intact neutrophils were observed prior to stimulation with *E. coli* with clearly defined boundaries for the DNA and G4 staining (Fig. 1, A-F). As expected, we observed minimal signal from the anti-citrullinated histone antibody in unstimulated neutrophils confirming that the isolated neutrophils were native and unstimulated. Stimulation with *E. coli* resulted in NETs as observed by the lack of defined cellular boundaries, formation of mesh-like structure in the DNA and G4 staining, and strong signal from the citrullinated histones (Fig. 1, G-L). Additionally, we observed a considerable degree of colocalization between G4 structures and hemin on the NETs, indicating the presence of stable G4 structures in the extruded DNA of NETs and suggesting a potential binding interaction between G4 and hemin. While intracellular G4/H interactions (*9–11*) have previously been described in other cell types, this co-localization offers evidence for the existence of extracellular G4/H in a biological setting.

**Fig. 1.**
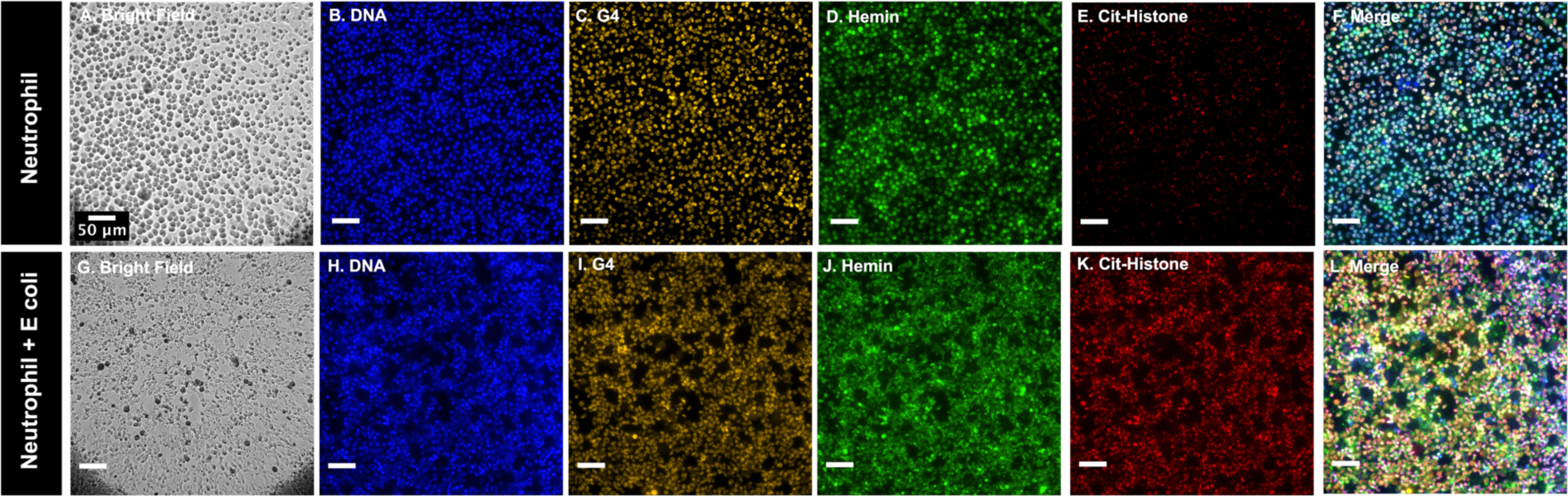
Immunofluorescence stain of G4/H colocalization on NETs. (**A-F**) Unstimulated neutrophils. (**A)** bright field. (**B**) condensed chromatin within nuclei, Hoechst 33342 stain. (**C**) G4 structures within nuclei, BG4 antibody staining. (**D**) 1D3 antibody staining of hemin. (**E**) anti-Citrullinated Histone H3 antibody staining of Histone H3. (**F**) Merge of B-E portraying defined cellular boundaries with intracellular labeling. (**G**-**L**) *E. coli* stimulated neutrophils. (**G**) bright field. (**H**) Extruded NETs DNA, Hoechst 33342 stain. (**I**) BG4 antibody staining of G4 structures within NETs. (**J**) 1D3 antibody staining of hemin with NETs. (**K**) anti-Citrullinated Histone H3 antibody staining of Histone H3 within NETs confirming the formation of NETs after *E. coli* stimulation. (**L**) Merge of GL showing extensive colocalization of G4 and hemin on NETs. Scale bar = 50 um.

Hemin is an important co-factor that is typically believed to be generated from the turnover of old red blood cells, and the extracellular levels of hemin can be exacerbated by hemolysis during infection. Surprisingly, we observed a strong hemin signal in both neutrophils from healthy donor and their extruded NETs without the addition of extraneous hemin in our experiment. Given that G4s in the genome are believed to sequester free hemin to protect from hemin-derived oxidative stress (*10*, *11*), it is possible that the hemin was sequestered by the G4s in neutrophils prior to the blood draw for the assay. However, since extracellular hemin can induce NETosis of neutrophils in a dose-dependent manner (*15–17*), it reduces the likelihood of the neutrophils in this study having come across free hemin prior to blood draw and raises the possibility of endogenously produced hemin in neutrophils.

### G4/H in NETs possess DNAzyme peroxidase activity

Previous *in vitro* studies have reported the peroxidase activity of G4/H complexes and that these synthetic DNAzymes have higher activity compared to that of just hemin alone (*12*, *14*, *18*). We sought to determine if the G4/H complexes in NETs possessed the same DNAzyme peroxidase activity to produce free radicals (FR). Isolated neutrophils form healthy donor were stimulated with phorbol myristate acetate (PMA) and the NETs DNA was then purified. We assessed the DNAzyme activity of purified NETs DNA to ensure we were only assaying the G4/H activity, excluding NETs associated proteins. To compare the activity against that of horseradish peroxidase (HRP) and commercial telomeric G4 complexed with hemin (Telo G4/H), we generated a standard curve of HRP activity on Amplex UltraRed (AR) and determined the amount of Telo G4/H that possessed the same activity (fig S1). For all subsequent experiments, we used the same mass of Telo G4 and purified NETs. The peroxidase activity of the individual G4 and hemin components was assessed, revealing no significant observable activity. However, in the presence of hydrogen peroxide (H_2_O_2_), when hemin was combined with NETs G4, the resulting G4/H complex exhibited 7.6 times higher peroxidase activity compared to hemin alone. Purified NETs demonstrated only 57% enzymatic activity compared to Telo G4/H despite the equal masses. This observation can be attributed to the likely lower abundance of G4 structures within the purified NETs or the presence of various topologies of G4 in NETs since non-parallel G4 have lower peroxidase activity (*19*–*21*). Notably, intriguing results were obtained when investigating the impact of various treatments on the DNAzyme activity. Surprisingly, the G4-specific competitive inhibitor BRACO19, the G4-binding hemin analog NMM, and the free radical scavenger vitamin C were found to abrogate the NETs DNAzyme activity by 92.5%, 83.5%, and 99%, respectively. In contrast, surprisingly, treatment with nucleases, including DNase I or EcoR I, did not result in a discernible reduction in enzymatic activity (Fig. 2.) suggesting that these nucleases were unable to degrade the non-canonical G4 structures (*22*, *23*). Although the DNAzyme exhibited lower activity compared to HRP, recent investigations have revealed its dependency on the specific substrate being catalyzed (*24*, *25*). Furthermore, with potentially >700,000 G4/H localized on NETs, greater quantities of FR with higher local concentrations could be produced.

**Fig. 2.**
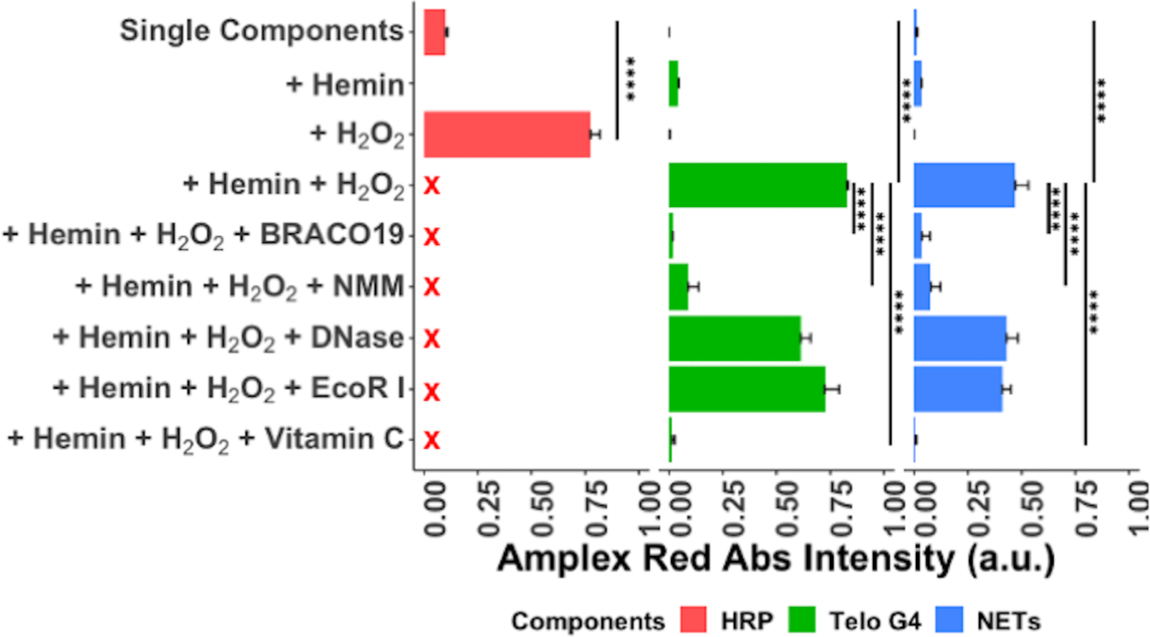
Peroxidase activity of G4/H DNAzyme in NETs using Amplex Red. Peroxidase activity using equal masses of Telo G4 and NETs were assayed with various combinations of individual components, G4 specific binding inhibitors (BRACO19, NMM), and antioxidant (Vitamin C). Telo G4+Hemin+H2O2 and HRP+H2O2 used have comparable activity. NETs+Hemin+H2O2 had greater activity than individual components and 7.6x higher than hemin alone. Peroxidase activity was abrogated by G4 inhibitors and antioxidant. All experiments were peformed in triplicate with mean and standard deviation plotted. Pairwise t-test with Bonferroni correction perfoemed to compare the different combinations. **** indicates p-value < 0.0001.

### G4/H DNAzyme in NETs produces FR and exerts a localized, concentrated effect on trapped bacterial cells

In order to ascertain the type of FR generated by the G4/H DNAzyme in NETs, their potential impact on bacteria, and the role of NETs DNA structure, we performed the proximity labeling assay on *Enterobacter cloacae* (EC). Proximity labeling exhibits optimal effectiveness within the 1-10 nm radius (*26*). We used biotin-phenoxyl radical, resulting from the generation of hydroxyl radicals (OH•) by the G4/H DNAzyme, to fluorescently label the surface of bacteria trapped within purified NETs. The generated biotin-phenoxyl radical selectively reacts with electron-rich amino acids, such as tyrosine, tryptophan, and others, located in close proximity on the bacterial surfaces. This reaction leads to fluorescence labeling upon treatment with fluorescein-streptavidin. EC was incubated with a biotin-phenol substrate, fluorescein-streptavidin, and a combination of purified NETs from PMA-stimulated neutrophils with NETs DNA of fragment size over 10 kilobases (Fig S2.) (*27*), hemin, and H_2_O_2_. Baseline fluorescence intensities from individual components were minimal. While the proximity labeling assay confirmed the production of OH• by the G4/H DNAzyme in NETs, the amount of labeling by OH• produced by the G4/H DNAzyme in NET was 137% and 160% greater than that by HRP and free-floating Telo G4 respectively (Fig. 3) despite being normalized according to activity in the standard curve described above suggesting an increased effect because of the structure of intact NETs. This was confirmed by digesting the NETs structure with EcoR I or DNase I, both of which reduced the labeling to 44% and 54% respectively, comparable to the Telo G4/H, likely because even short oligonucleotides can result in DNA-protein interactions on the cell surface (*28*, *29*). These findings support the local concentrated effects of the G4/H DNAzyme on trapped bacteria in NETs, especially since mass for mass, we observed earlier that Telo G4/H has greater peroxidase activity than purified NETs G4/H DNAzyme. In addition, while monomeric G4/H DNAzyme units have lower activity than peroxidase enzymes like HRP, it is possible that G4/H DNAzyme in bacteria-bound NETs formed multimeric units that have increased synergistic activity (*30*). As expected, the trapping effect was not readily apparent for *Staphylococcus aureus* (SA) (Fig. S3) since it is known to evade NETs through its ability release nucleases to breakdown DNA (*31*) resulting in comparable signal between free-floating Telo G4, intact NETs, and additional EcoR I or DNase I treatments. Interestingly, although NETs are known to produce various FR like HOCl and NO• through the action of myeloperoxidase and nitric oxide synthase respectively, no enzymatic generators of OH• in NETs were previously known. Our evidence suggests that not an enzyme but the G4/H DNAzyme produces OH• in NETs.

**Fig. 3.**
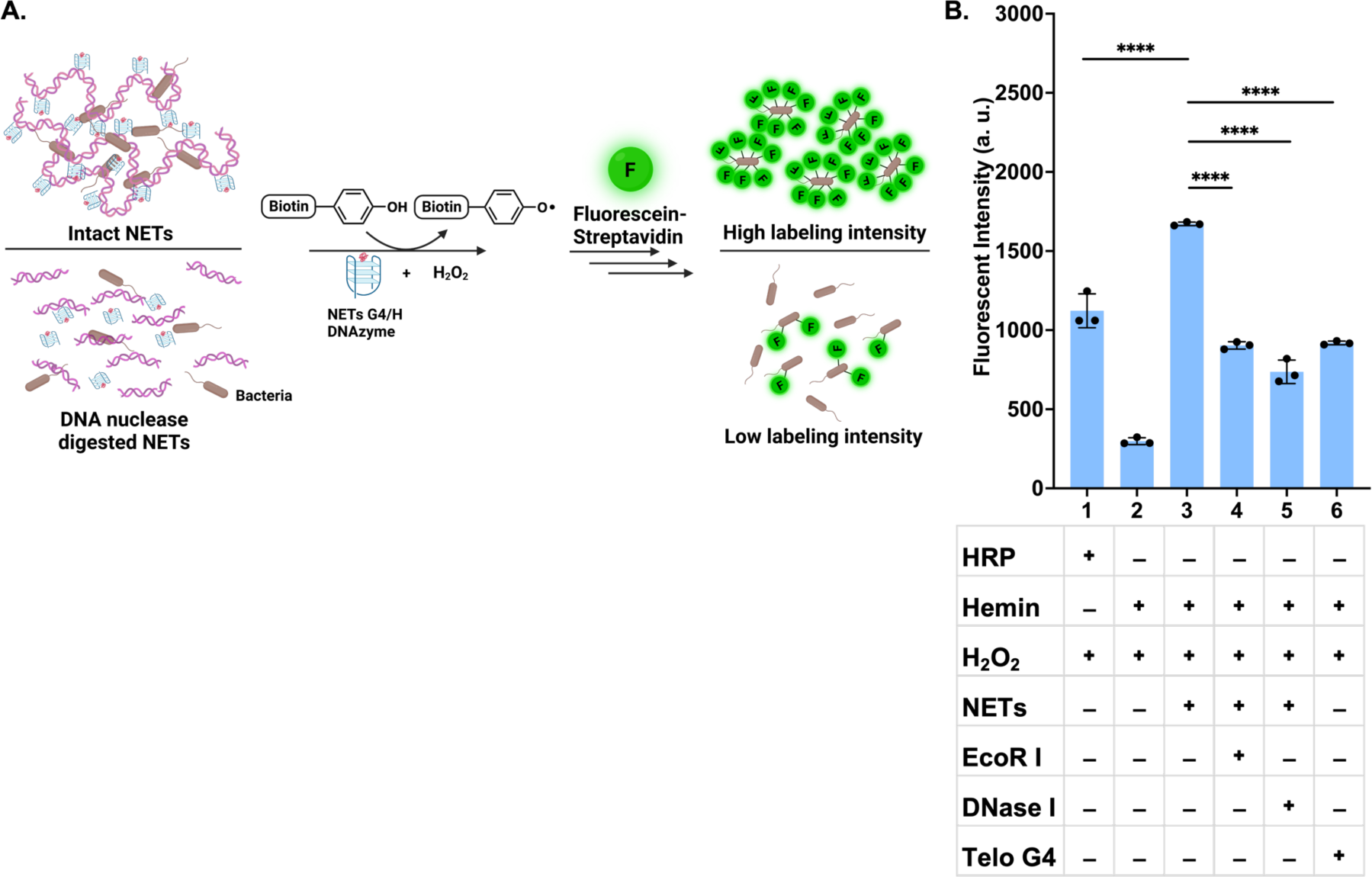
NETs DNA-based proximity labeling assay. **(A**) Schematic of proximity labeling assay. Labeling of bacteria trapped in NETs, versus untrapped bacteria in nuclease-digested NETs, with biotin-phenoxyl radicals by the DNAzyme activity of G4/H complexes. (**B**) Purified NETs from PMA-stimulated neutrophils incubated with EC, biotinphenol, hemin, H2O2, and fluorescein-streptavidin to allow for bacterial surface labeling. Upon digestion of NETs with EcoR I and DNAse I on the same bacteria, the resulting fluorescent intensity showed a reduced labeling effect comparable to that of free-floating Telo G4 (∼54%) compared to intact NETs. (n = 3 biologically independent experiments; bars represent mean signal, and error bars denote s.e.m.; one-way ANOVA performed; **** indicates p-value < 0.0001)

Since OH•, the most reactive FR *in vivo* (*32*), is known to have bactericidal activity, we assessed the *in vitro* bactericidal activity of the G4/H DNAzyme in purified NETs using a plating assay against EC (Fig. S4). EC was incubated with different components of the G4/H DNAzyme individually or in combination along with H_2_O_2_, and the extent of bacterial killing was assayed. Neither hemin nor NETs G4 alone was able to kill EC after overnight incubation. Additionally, 5mM H_2_O_2_ did not result in significant bacterial killing, likely because EC encodes the catalase enzyme that breaks down H_2_O_2_ to water and oxygen. In line with expectations, the G4/H DNAzyme in NETs achieved a 94% reduction in EC viability with a significant decrease in EC within 20 minutes (Fig. S5) suggesting potent bactericidal activity of the G4/H DNAzyme in NETs. DNA, the largest component of NETs was only believed previously to be a scaffold for associated proteins and to trap invading pathogens with potential for bactericidal activity through chelating properties of the DNA backbone (*1*, *5*). Our *in vitro* assays suggest a potent role for the G4, a non-canonical DNA secondary structure of NETs in clearing pathogens.

### G4/H DNAzyme is physiologically active and the major contributor of bactericidal activity in NETs

We next sought to determine if the G4/H DNAzyme was physiologically active and measure its contribution to the bactericidal activity of NETs *ex vivo.* We hypothesized that all essential components for its activity were naturally present in blood, for example during inflammation, hemolysis increases extracellular hemin levels, and dissolved H_2_O_2_ in plasma can be as high as 50 μM (*33*, *34*). To demonstrate this phenomenon, we conducted an *ex vivo* experiment with neutrophils isolated from a healthy individual against EC, following a previously published protocol with minor modifications (*1*). EC was incubated with neutrophils in DPBS containing heat-inactivated human serum along with various combinations of interleukin 8 (IL-8), cytochalasin D (an antiphagocytic agent), BRACO19, NMM, and DNase I with no additional hemin or H_2_O_2_. Stimulation of neutrophils with IL-8 led to approximately 85% eradication of EC (Fig. 4) despite the ability of EC to encode catalase to protect it from H_2_O_2_ toxicity likely because synthetic DNAzymes have a higher affinity for H_2_O_2_ than catalases (*35*, *36*). Interestingly, inhibiting phagocytosis had only a marginal impact on the bactericidal efficacy, indicating that the majority of the observed killing in our assay can be attributed to NETs. Administration of NMM and BRACO19, which specifically target the G4/H structure of NETs, resulted in a substantial reduction of the bactericidal effect to less than 20%. Interestingly, quenching FR through treatment with vitamin C resulted in less than 20% bactericidal effect similar to treatment with BRACO19 and NMM suggesting that in our *ex vivo* model, the other mechanisms of bacterial killing in NETs contributed minimally and the majority of the FR was generated by the G4/H DNAzyme. Similar results were observed with SA (Fig. S6), however, with less overall killing than with EC likely because it encodes extracellular nucleases. Remarkably, treatment with DNase I exhibited only a modest decrease of ∼20% in the bactericidal activity against EC but ∼80% decrease in killing of SA similar to findings from previous studies (*1*, *5*). Notably, DNase I does not disrupt the G4 structure (*22*, *23*) or its peroxidase activity, as demonstrated in Fig. 4. Therefore, the observed disparity can be attributed to the elimination of bacterial entrapment by digested NETs that affects different bacteria to varying degrees, while the bactericidal effect is still retained through the intact G4/H DNAzyme activity. This *ex vivo* investigation not only confirms the physiological occurrence of G4/H DNAzyme activity within NETs, but also highlights the prominent role of the G4/H DNAzyme as the primary contributor to FR generation and subsequent bactericidal activity exhibited by NETs.

**Fig. 4.**
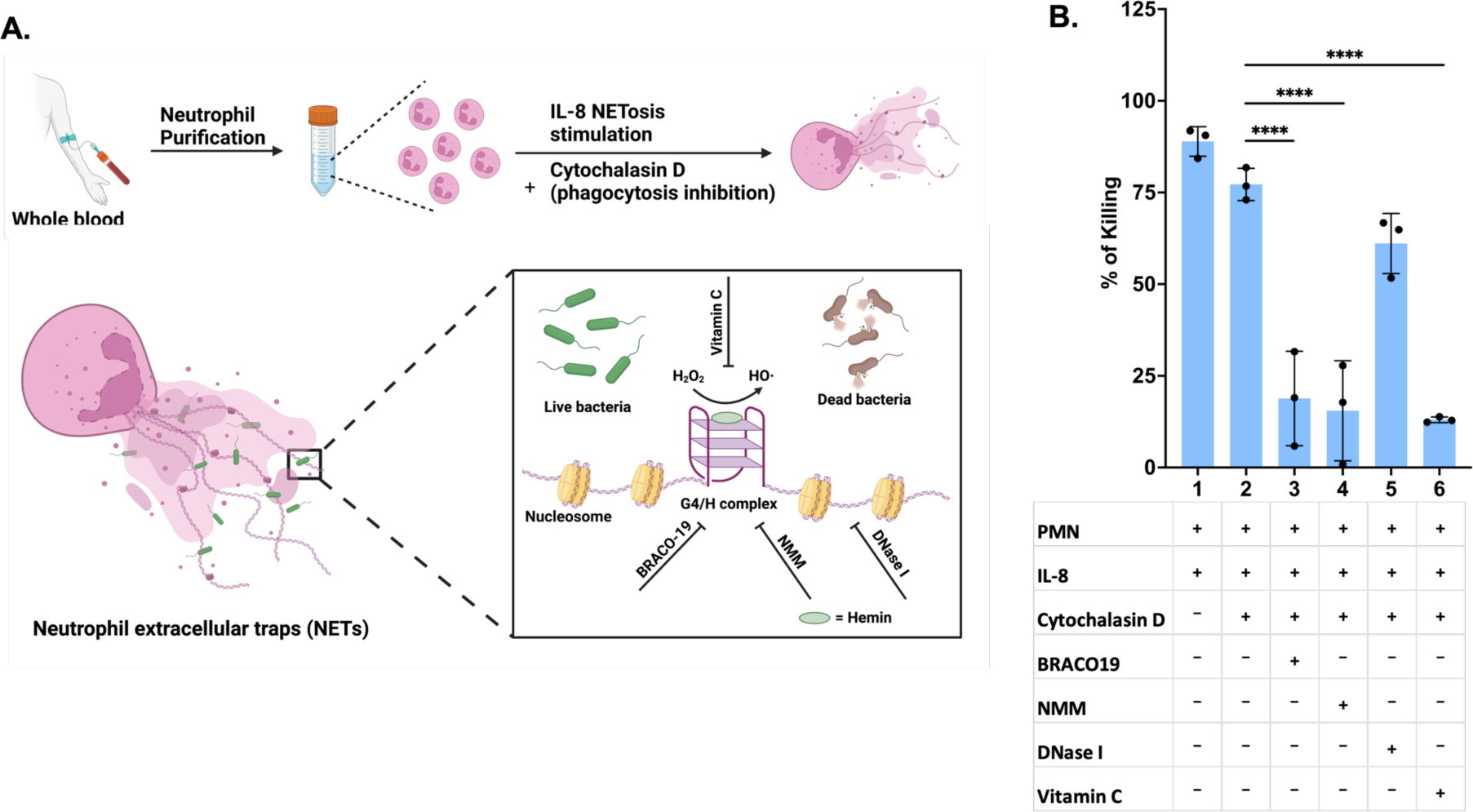
*Ex vivo* bactericidal activity of the G4/H DNAzyme against EC. (**A**) Schematic of the *ex vivo* bactericidal assay. Neutrophils from whole blood were stimulated with IL-8 and treated with various combinations of antiphagocytic cytochalasin D, G4 specific inhibitor BRACO19, G4 binding hemin analog NMM, and DNase I with no additional hemin or H2O2. (**B**) Bactericidal activity of isolated neutrophils through NETs against EC. ∼ 80% of inoculated EC was killed by IL-8-stimulated NETs. Phagocytosis account for an additional ∼10% of killing. Abrogation of NETs killing by G4-specific inhibitors like BRACO19, NMM, or antioxidant Vitamin C (< 20%). DNase I treatment only reduced killing by ∼20%. (n = 3 biologically independent experiments; bars represent mean signal, and error bars denote s.e.m.; one-way ANOVA performed; **** indicates p-value < 0.0001)

Synthetic G4/H DNAzyme has been studied for years with various applications in biosensors and other biochemical assays (*14*, *37*, *38*), but their physiological existence and biological functions are not widely understood. G4 structures in the genomic landscape have been detected to sequester free hemin both by DNA and RNA forms to protect the cell from oxidative damage and affect transcriptional control (*9*, *11*). However, these studies did not describe the peroxidase activity of the biological G4/H complex. In this study, we highlight the existence and biological activity of the G4/H DNAzyme in the extruded, decondensed chromatin present in NETs. The G4/H DNAzyme mimics peroxidase activity in producing OH• from H_2_O_2_ and has stronger activity than hemin alone. Trapping of bacteria in NETs DNA subjects it to increased local concentrations of the OH• resulting in the mechanism being the driver of bactericidal activity in NETs. While our demonstration is in an extracellular setting, our findings raise the question of the effect of highly localized OH• on the genome intracellularly, especially when hemin is sequestered. It is possible that one of the hundreds of putative G4 interacting proteins (*39*) recently discovered dampens the DNAzyme activity within the cell, but further work is needed to elucidate these mechanisms. Additionally, we only assayed the role of the G4/H generated FR in bactericidal activity associated with NETs while there may be other biological consequences. For example, FRs from NETs have been associated with host tissue damage and the formation of posttranslational modifications on NETs-associated histones resulting in autoimmune disorders like systemic lupus erythematosus (*40–44*). We offer a paradigm shift in the current understanding of the role of the NETs DNA – beyond a scaffold to the driver of potentially multiple biological consequences of NETs.

## Acknowledgments

We would like to thank Alice Y Ting for advice with assays and a critical review of the manuscript.

## Author contributions

Conceptualization: TK, SY

Methodology: TK, SY

Investigation: TK, EJZ

Visualization: TK, NR

Funding acquisition: SY

Project administration: SY

Supervision: SY

Writing – original draft: NR, TK, SY

Writing – review & editing: NR, TK, EJZ, RA, SY

## Data and materials availability

All data are available in the main text or the supplementary materials.

## Supplementary Materials

### Materials and Methods

#### Isolation of human blood neutrophils

Human neutrophils were isolated from a healthy donor using the EasySep Direct Human Neutrophil Isolation Kit, following the protocol provided by the vendor (#19666, StemCell Tech, Vancouver, BC, Canada). The isolated neutrophils were then resuspended in DPBS (#14190-144, Gibco, ThermoFisher) without calcium and magnesium, supplemented with 1 mM EDTA (#AAJ15694AE, Fisher Scientific).

#### *In vitro* neutrophil NETosis stimulation, NETs isolation and quantification

The freshly isolated neutrophils were immediately seeded in 12-well plates at a density of 1×106 cells per well. The cells were incubated at 37°C for 30 minutes and then stimulated with 100 nM phorbol 12-myristate 13-acetate (PMA) (#400145, Cayman Chemical) in phenol-red free RPMI-1640 (#11835030, Gibco, ThermoFisher Scientific) supplemented with 2 mM calcium chloride (#21115, MilliporeSigma) at 37°C in the presence of 5% CO2. After 4 hours, each well was carefully washed twice with 1 mL of DPBS and then treated with 500 μl of DPBS containing 10U Alu I restriction enzyme (#ER0011, Thermo Scientific) for 30 minutes at 37°C. The supernatant from each well was collected and centrifuged for 5 minutes at 300 x g at 4°C to remove cells and cell debris. The supernatants rich in NETs were then further purified using the Monarch genomic DNA purification kit (#T3010L, New England Biolabs). The purified NETs DNA was preserved in sterilized DNase-free water and quantified using the Quant-iT PicoGreen dsDNA Kit (#P11496, Molecular Probes, ThermoFisher Scientific), following the manufacturer’s instructions.

#### Immunofluorescence staining for NETs visualization

We modified Biffi et al (*45*) human cell G-quadruplexes immunofluorescence staining method for NETs G-quadruplex visualization. Briefly, we isolated human neutrophils from a healthy donor. The freshly isolated neutrophils (5×10^5^) were immediately seeded in 4-well chamber slides (#154526, ThermoFisher) with 1ml phenol-red free RPMI-1640 supplemented with 2 mM calcium chloride ((#21115, MilliporeSigma). The chamber slides were incubated in a CO2 incubator under 37 °C for 20 minutes. The neutrophils were subjected to treatment with or without Escherichia coli for a duration of 2 hours to initiate NETosis and generate NETs. The generated NETs were rinsed twice with cold DPBS and then fixed with 4% paraformaldehyde for 30 minutes at room temperature. Subsequently, the NETs were washed three times with DPBS and permeabilized with 0.1% Triton X-100 in DPBS for 1 hour at 4 °C. Afterward, the NETs were washed three more times with DPBS and treated with 50 μg/ml RNase A (#T3018, New England Biolabs) for 1 hour at 37 °C. Following this, the NETs were blocked with a mixture of 1% skim milk and 0.1% Tween-20 in DPBS at 4 °C overnight. Immunofluorescence stain was performed using standard methods with BG4 (#MABE917, MilliporeSigma), anti-FLAG (#F1804, MilliporeSigma) and anti-mouse IgG Alexa 594 (#AB150116, abcam) antibodies for detection G-quadruplex; 1D3 (#Ab00982-10.0, absolute antibody) and anti-human IgG Alexa-488(#A-11013, ThermoFisher Scientific) antibodies for Hemin; anti-Cit-Histone (ab5103, abcam) and anti-rabbit IgG Alexa 647 (ab150083, abcam) antibodies for citrullinated-Histone. NETs DNA was stained with Hoechst 33342 (62249, ThermoFisher Scientific). Digital images were recorded using a Nikon ECLIPSE Ti2-E microscope and analyzed with Image J software.

#### *In vitro* bactericidal assay

Monoclonal bacteria, *Enterobacter cloacae*, were cultured in LB medium and shaken at 37 °C overnight before being used. The bacterial cells were collected by centrifugation and then resuspended in a 10 mM KCl buffer (#60142, MilliporeSigma) to achieve an optical density of 1.0 at 600 nm (OD 600). Subsequently, the bacteria were diluted to 0.03 OD 600 using a sterile KCl buffer. The prepared bacterial solution was then combined with various combinations of 300 ng NETs DNA, 300 ng Telo G4, 10uM Hemin, and 5 mM H_2_O_2_, and incubated at 37 °C for 30 minutes. Afterward, the solution was diluted 10 times, and an aliquot of 50 μL of the bacterial suspension was spread onto an NB agar plate for overnight growth in a 37 °C incubator. The bacterial colonies on the plates were then counted. Telo G4 sequence: TTAGGG TTAGGG TTAGGG TTAGGG TTAGGG TTAGGG TTAGGG TTAGGG TT (Integrated DNA Technologies).

#### NETs DNA fragmentation and electrophoresis

300 ng purified NETs DNA was subjected to distinct enzymatic treatment using 20 units EcoR I (#R0101S, New England Biolabs) or 4 units DNase I (#M0303L, New England Biolabs) for 60 minutes at 37°C, respectively. Subsequently, both enzymatic reactions were terminated through heat inactivation at 75°C for 10 minutes. The resulting NETs DNA samples were subjected to electrophoresis on a 1.3% agarose gel, followed by visualization using ethidium bromide staining.

#### NETs DNAzyme peroxidase assay

In this study, we investigated the peroxidase capability of the G4/H DNAzyme within purified NETs DNA induced by PMA. Specifically, we assessed its capacity to oxidize Amplex UltraRed (AR) reagent (#A36006, ThermoFisher Scientific) in the presence of hydrogen peroxide (H_2_O_2_). ((#A22188, Component E, ThermoFisher Scientific) A standard curve of horseradish peroxidase (HRP) (#A22188, Component D, ThermoFisher Scientific) activity on AR was initially established to serve as a reference. Subsequently, we determined the optimal quantity of Telo G4 (300 ng, equals 2 mU of HRP) required to achieve comparable enzymatic activity. The same mass of NETs DNA was employed as Telo G4. 4 units DNase I, 20 units EcoR I, 150 uM G4-specific competitive inhibitor BRACO19 (#SML0560, MilliporeSigma), the G4-binding hemin analog 150 uM N-methyl mesoporphyrin IX (NMM) (#396879, Santa Cruz Biotechnology), and 2 mM free radical scavenger vitamin C (#11140, MilliporeSigma) were tested to abrogate the DNAzyme activity.

#### NETs DNA proximity labeling assay

Single colonies of *Enterobacter cloacae* and *Staphylococcus aureus* were cultured in the LB medium and shaken at 37 °C overnight before usage. The bacterial cells were collected by centrifuging, DPBS washing 5 times and redispersed in 10 ml KCl buffer with an optical density of 1.0 at 600 nm (O.D.600). 1 ml bacterial solution was mixed with different combinations of 6.67 ug NETs DNA, 3 ug Telo G4, 10 uM Hemin, 10 mM H_2_O_2_ and 10 mM Biotin-tyramide (#SML2135, MilliporeSigma) reagent incubating at 25 °C for 45 minutes. After being washed five times with DPBS, the sample was incubated with FITC Conjugated Avidin (10ug/ml) (#21221, ThermoFisher Scientific) for a duration of 2 hours. Following this incubation, the sample is washed five more times with DPBS before measuring the fluorescence signals using a fluorescent plate reader (BioTek Synergy LX, Agilent).

#### *Ex vivo* bactericidal assay

Neutrophils were collected using the previously described method. The freshly isolated neutrophils were promptly seeded into 12-well plates at a density of 2×10^6^ cells per well in 500 μl DPBS containing 10 ng IL-8 and 2 mM CaCl_2_ and then agitated at 37 °C for 20 minutes. Subsequently, 10 μl of heat-inactivated human serum with cytochalasin D (10 μl/ml) (#C8273, MilliporeSigma) was added to each well, followed by shaking and incubation for 15 minutes. Next, the samples were treated with 100 U/ml of RNase for 10 minutes. To demonstrate the significance of NETs DNAzyme in extracellular killing, 50 uM BRACO19, 50 uM NMM, 10U DNAase I, and 2 mM Vitamin C were pre-treated 5 minutes prior to bacterial addition (5×10^3^ bacteria/well). The samples were then agitated and incubated for 40 minutes. Afterward, each sample was cultured on Nutrient Agar Plate overnight for colony counting.

## Supplementary figures

**Fig. S1.**
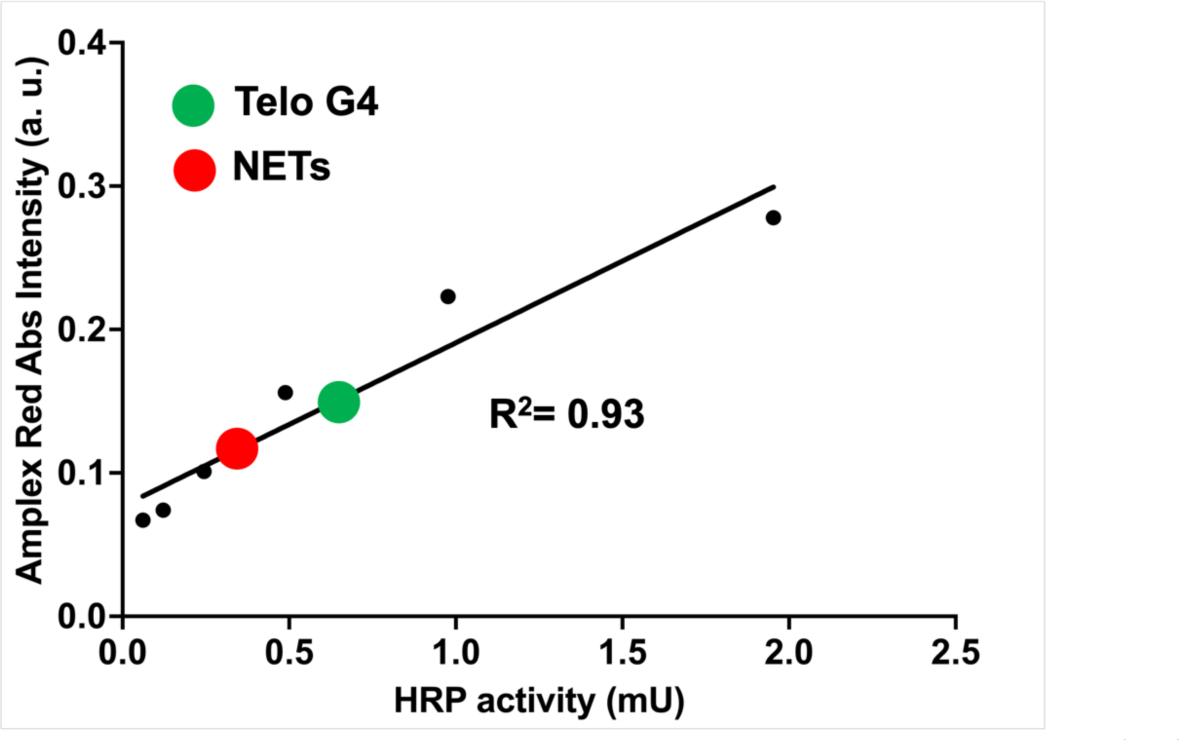
Estimating amount of Telo G4 needed to achieve same HRP activity. HRP standard curve was stablished by using Amplex UltraRed as substrate. 100 ng of NETs DNA and Telo G4 was used to determine the enzymatic activity.

**Fig. S2.**
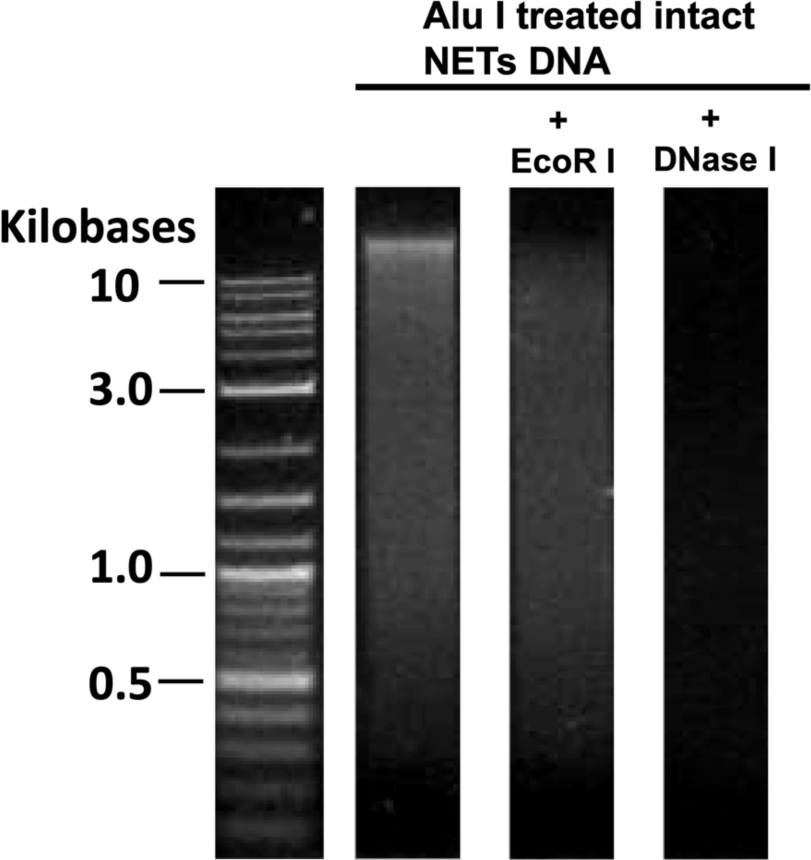
Purified NETs DNA fragmentation. Purified NETs DNA gel electrophoresis after Alu I digestion and genomic DNA purification with or without further EcoR I, DNase I treatment. DNA migration took place in 1.3% agarose gel staining with ethidium bromide.

**Fig S3.**
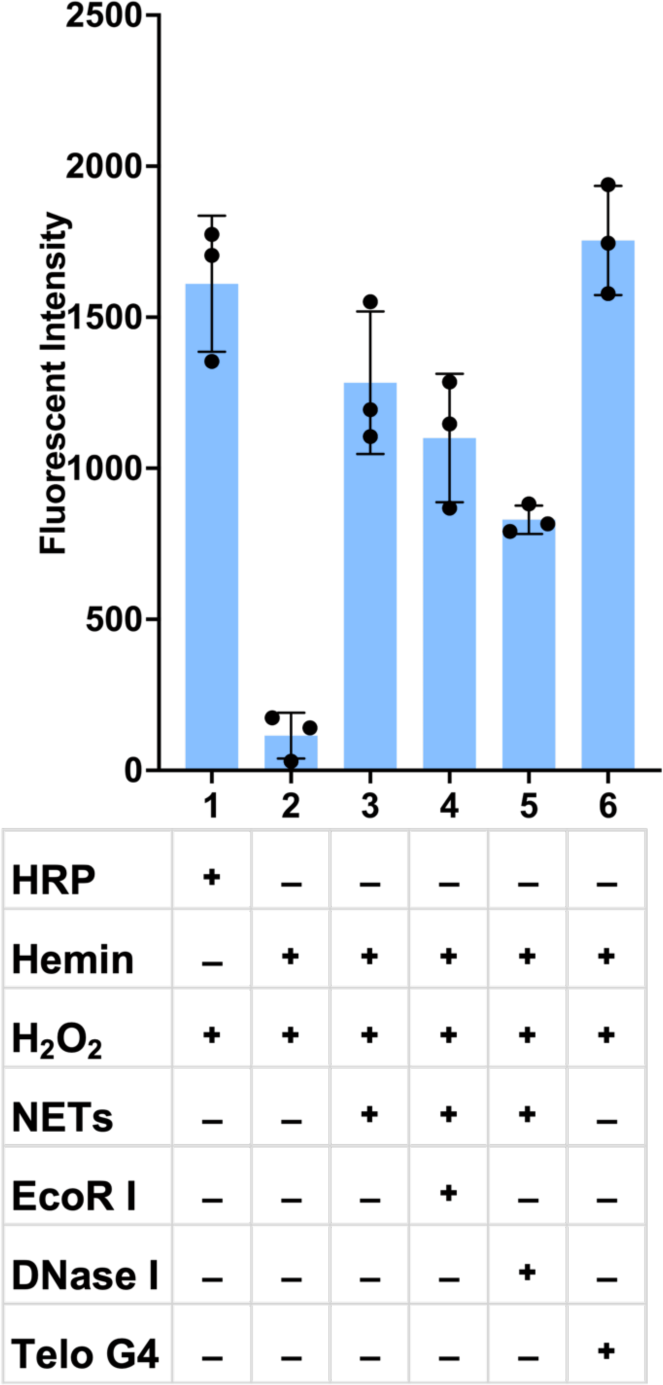
Proximity labeling of *Staphylococcus aureus* (SA). Purified NETs from PMA-stimulated neutrophils incubated with SA, biotin-phenol, hemin, H2O2, and fluorescein-streptavidin to allow for bacterial surface labeling. Upon digestion of NETs with EcoR I and DNAse I on the same bacteria, the resulting fluorescent intensity showed a reduced labeling effect compared to intact NETs. (n = 3 biologically independent experiments; bars represent mean signal, and error bars denote s.e.m. one-way ANOVA performed; no significant difference was found between HRP, NETs, Telo G4, DNase and EcoR I treatment.)

**Fig. S4.**
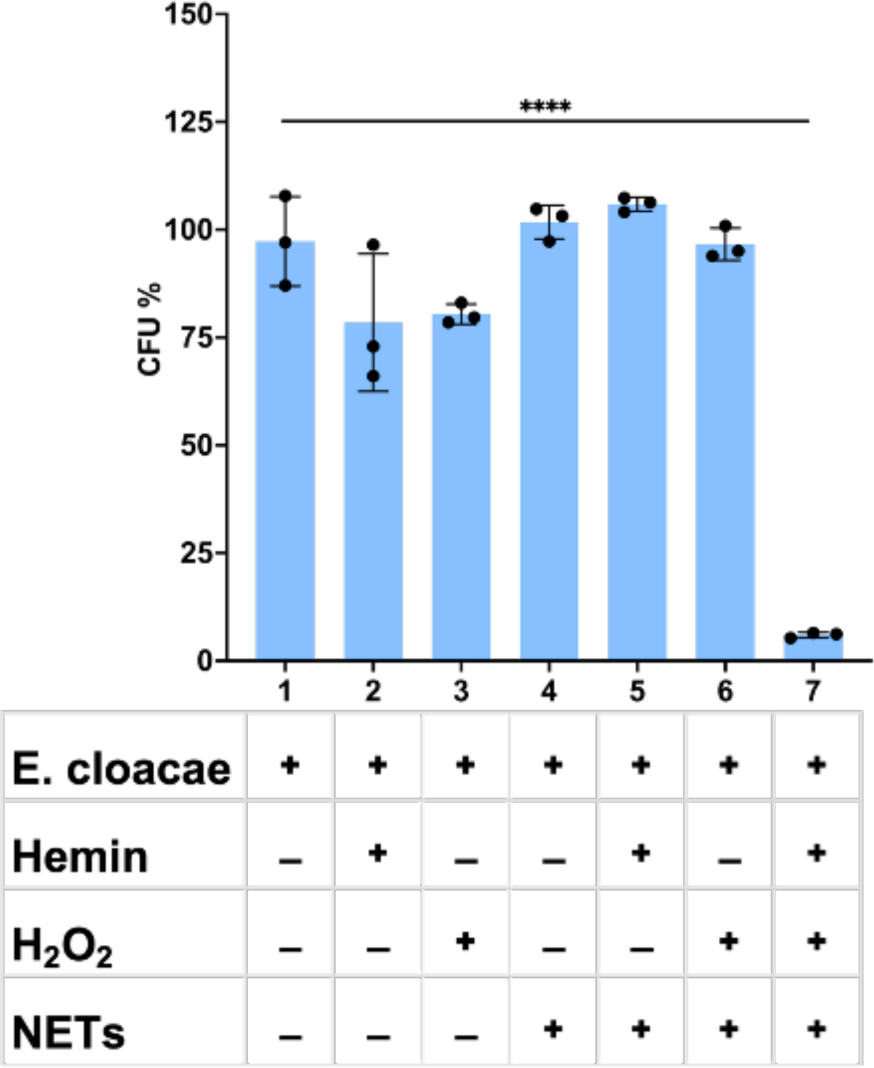
*In vitro* bactericidal assay. EC was incubated with various combinations of purified NETs from PMA-stimulated neutrophils, hemin, and H2O2 to allow for bacterial killing. Plate colony counting was performed after overnight incubation. Individual components were unable to kill EC. Only complete DNAzyme with H2O2 killed EC (n = 3 biologically independent experiments; bars represent mean signal, and error bars denote s.e.m. one-way ANOVA performed; **** indicates p-value < 0.0001)

**Fig. S5.**
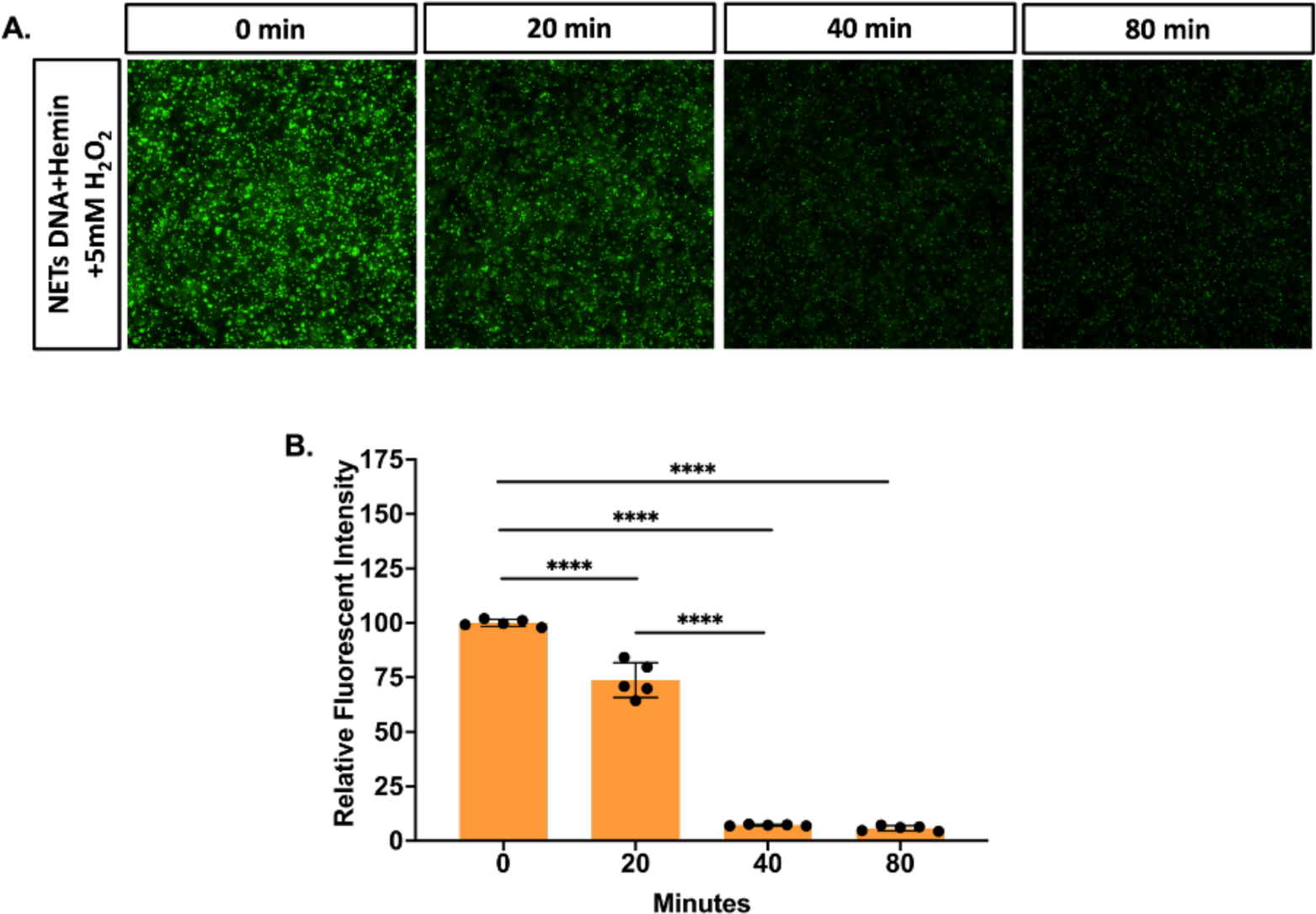
Time-dependent bactericidal assay. *E. coli* was incubated with purified NETs DNA, hemin, H2O2, and SYTO9 dye to allow for monitoring bacterial viability. (**A**)SYTO 9 fluorescent image of living bacteria at various time points. (**B**) Image fluorescence intensity at each time points. (n = 5 biologically independent random field; bars represent mean signal, and error bars denote s.e.m. one-way ANOVA performed; **** indicates p-value < 0.0001)

**Fig. S6.**
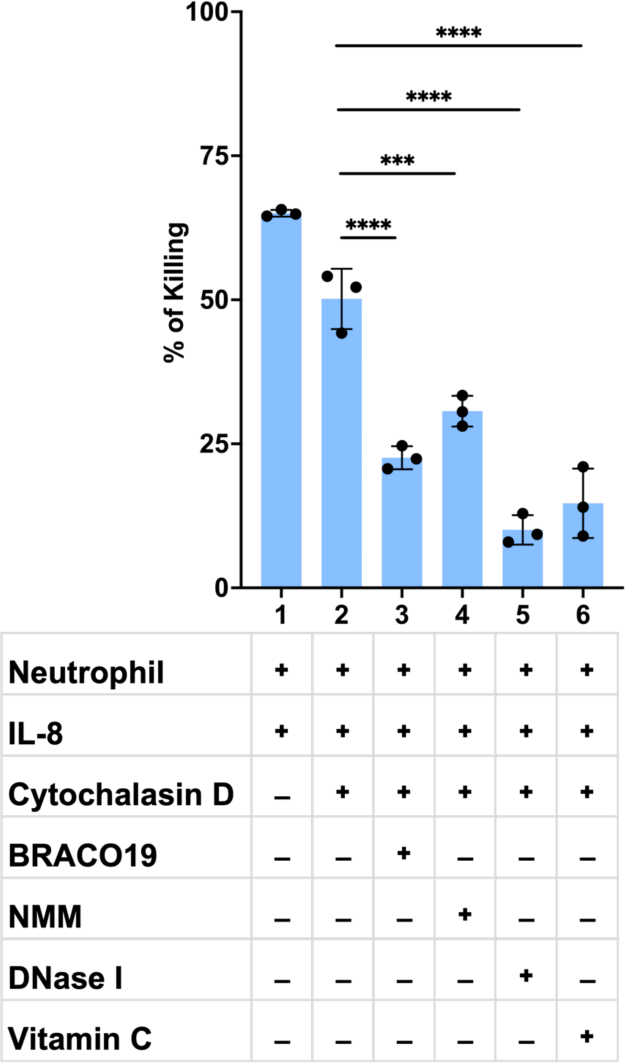
*Ex vivo* bactericidal activity of the G4/H DNAzyme for *Staphylococcus aureus* (SA). Bactericidal activity of isolated neutrophils through NETs. ∼ 63% of inoculated SA was killed by IL-8-stimulated NETs. Phagocytosis account for an additional ∼13% of killing. Abrogation of NETs killing by G4-specific inhibitors like BRACO19, NMM, or antioxidant Vitamin C (< 25%). (n = 3 biologically independent experiments; bars represent mean signal, and error bars denote s.e.m.; one-way ANOVA performed; **** indicates p-value < 0.0001, *** indicates p-value < 0.001)w

